# Comparative Tn-Seq reveals common daptomycin resistance determinants in *Staphylococcus aureus* despite strain-dependent differences in essentiality of shared cell envelope genes

**DOI:** 10.1101/648246

**Authors:** Kathryn A. Coe, Wonsik Lee, Gloria Komazin-Meredith, Timothy C. Meredith, Yonatan H. Grad, Suzanne Walker

## Abstract

Antibiotic-resistant *Staphylococcus aureus* remains a leading cause of antibiotic resistance-associated mortality in the United States. Given the reality of multi-drug resistant infections, it is imperative that we establish and maintain a pipeline of new compounds to replace or supplement our current antibiotics. A first step towards this goal is to prioritize targets by identifying the genes most consistently required for survival across the *S. aureus* phylogeny. Here we report the first direct comparison of gene essentiality across multiple strains of *S. aureus* via transposon sequencing. We show that mutant fitness varies by strain in key pathways, underscoring the importance of using more than one strain to differentiate between core and strain-dependent essential genes. Despite baseline differences in gene importance, several pathways, including the lipoteichoic acid pathway, become consistently essential under daptomycin exposure, suggesting core vulnerabilities that can be exploited to resensitize daptomycin-nonsusceptible isolates. We also demonstrate the merit of using transposons with outward-facing promoters capable of overexpressing nearby genes for identifying clinically-relevant gain-of-function resistance mechanisms. Together, the daptomycin vulnerabilities and resistance mechanisms support a mode of action with wide-ranging effects on the cell envelope and cell division. This work adds to a growing body of literature demonstrating the nuanced insights gained by comparing Tn-Seq results across multiple bacterial strains.

**Author summary:** Antibiotic-resistant *Staphylococcus aureus* kills thousands of people every year in the United States alone. To stay ahead of the looming threat of multidrug-resistant infections, we must continue to develop new antibiotics and find ways of making our current repertoire of antibiotics more effective, including by finding pairs of compounds that perform best when administered together. In the age of next-generation sequencing, we can now use transposon sequencing to find potential targets for new antibiotics on a genome-wide scale, identified as either essential genes or genes that become essential in the presence of an antibiotic. In this work, we created a compendium of genes that are essential across a range of *S. aureus* strains, as well as those that are essential in the presence of the antibiotic daptomycin. The results will be a resource for researchers working to develop the next generation of antibiotic therapies.

## Introduction

Every year in the United States over two million people acquire an antibiotic-resistant infection, and over 23,000 people die from those infections [1]. More people die of infections caused by antibiotic-resistant *Staphylococcus aureus* than any other resistant pathogen [2]. The primary antibiotic class used to treat *S. aureus* infections is the β-lactams, but over 50% of *S. aureus* infections in the United States are resistant to methicillin and other β-lactamase-impervious β-lactams [3, 4]. These β-lactam-resistant strains are termed methicillin-resistant *S. aureus* (MRSA). Antibiotics such as vancomycin, linezolid, and daptomycin are used to treat MRSA infections but are ineffective in an increasing number of cases [5–7]. We need a pipeline of new compounds that can either kill *S. aureus* outright or resensitize antibiotic-resistant isolates, regardless of genetic background. Thousands of *S. aureus* isolates have been sequenced, revealing substantial differences in gene content [8, 9], but the functional variability across the phylogeny has yet to be systematically investigated. Transposon sequencing (Tn-Seq) can be used to attribute phenotypes to genes on a genome-wide scale, enabling researchers to characterize the full essential genome of a bacterial strain in a single experiment [10, 11]. Here we describe the first multi-strain Tn-Seq study in *S. aureus*, using comparisons of results from favorable growth conditions to define baseline variation in genetic dependencies and then exposing the libraries to daptomycin to probe the factors that modulate daptomycin susceptibility.

Several transposon mutagenesis studies have formerly analyzed the essential genes in individual *S. aureus* strains [12–17]. Aside from one study conducted in a livestock-associated strain, all the studies involved one clonal complex, CC8, and most emphasized laboratory-adapted strains. Only two methicillin-resistant strains have been examined. Moreover, the conditions for library creation and analysis differed substantially. With these limitations, it is difficult to assess the congruence of gene essentiality across *S. aureus* strains. A recent comparative Tn-Seq analysis of *Mycobacterium tuberculosis* showed strain-to-strain differences in gene essentiality, with implications for antibiotic targets and acquisition of resistance; this work highlighted the importance of carrying out functional genomics studies of important pathogens in multiple strains using the same methodology [18].

Here we have used our previously reported platform for creating transposon libraries in *S. aureus* to make high-density transposon libraries in five strains [17]. These libraries comprise six sub-libraries, each made using a different transposon construct. Several constructs contain outward-facing promoters, which minimize polar effects within polycistronic operons. In addition, under some conditions, insertions of outward-facing promoter constructs can upregulate nearby genes to provide a distinct fitness advantage that is mechanistically informative [19]. The five *S. aureus* strains we chose represent three clonal complexes (Fig 1B), including two MRSA/methicillin-sensitive (MSSA) pairs from the same sequence type. MW2, from sequence type 1 (ST1), was the first lineage of community-acquired MRSA infections reported in the United States [20–22]. MSSA476, a representative of a United Kingdom community-acquired MSSA lineage, is likewise from ST1 [23]. We have chosen USA300-TCH1516 to represent the hypervirulent USA300 lineage from ST8, which has become the most common community-acquired MRSA lineage in the United States [21, 24, 25]. The commonly used MSSA lab strain HG003 is also ST8. To capture a larger scope of the *S. aureus* phylogeny, we also included MRSA252, an ST36 hospital-acquired MRSA strain predominantly found in Europe that has attracted attention for its unusually large accessory genome [23].

**Fig 1.**
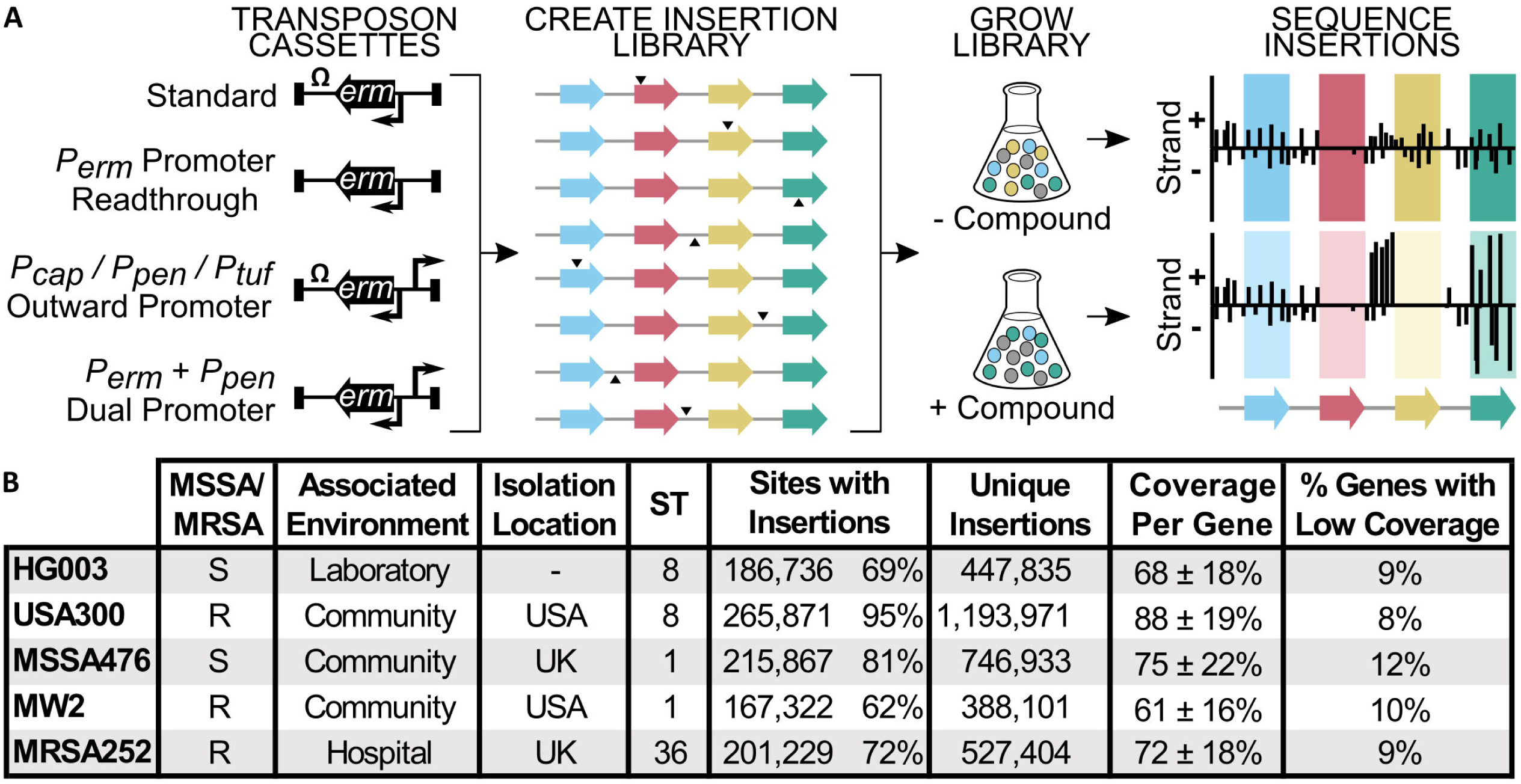
We created complex transposon libraries of similar coverage in five diverse *S. aureus* strains. (A) Six barcoded transposon constructs were used to make sub-libraries that were combined into a single transposon library. Construct 1 can drive expression only of *erm^R^* (gene within the transposon constructs). Constructs 2-6 can drive expression of other genes, either by readthrough of *erm^R^* in constructs lacking terminators or because they contain an additional outward-facing promoter (*P_cap_, P_pen_*, or *P_tuf_*). The libraries can be sequenced to find essential genes that lack transposon insertions (red gene) or treated with compound and then sequenced to find conditionally essential genes (yellow gene), upregulation signatures (upstream of yellow gene), and genes with conditional fitness costs (green gene). (B) High-density transposon libraries were made in five *S. aureus* strains. MSSA = methicillin-sensitive *S. aureus;* MRSA = methicillin-resistant *S. aureus*; ST = sequence type. Sites with insertions refers to the number of TA dinucleotide sites having insertions of at least one transposon construct. The “unique insertions” column sums the TA insertion counts from each transposon construct. Coverage per gene is the average percent of TA sites within a gene with insertions, ± standard deviation. Low-coverage genes were defined as those that had less than half of the average gene coverage.

We found 200 genes that are essential in all five strains and thus represent the core essential genome. We also identified genes that were uniquely essential in a given strain, as well as genes and pathways present in all five strains but essential in a subset. These cases imply that strain-specific distal genetic determinants, whether allelic in nature or horizontally acquired as with β-lactam resistance cassettes, can profoundly reshape the role of even core pathways. The lipoteichoic acid (LTA) pathway is one example of a conserved pathway that is differentially important across *S. aureus* strains.

We probed to what extent the variable genetic dependencies evident in the essential gene analysis shaped mutant fitness upon exposure to daptomycin, an antibiotic used for *S. aureus* infections. Patients who cannot tolerate β-lactam or vancomycin treatment, or who have resistant infections, are commonly treated with daptomycin. Although still rare, daptomycin nonsusceptibility is on the rise [6]. It might be possible to overcome infections that are resistant to daptomycin by co-administering a compound that targets an intrinsic resistance factor found to be universally important in withstanding daptomycin stress. A recent study comparing two *Streptococcus pneumoniae* strains found major differences in factors that influenced daptomycin susceptibility, raising questions about the prevalence of universally conserved daptomycin intrinsic resistance factors in *S. aureus* [26]. Nevertheless, we identified daptomycin intrinsic resistance factors conserved in all strains and demonstrated that even pathways that appear variably essential among strains under normal growth conditions can become essential in all upon exposure to daptomycin, illuminating a path forward for combatting daptomycin resistance in *S. aureus*.

## Results

### Creation of high-density *S. aureus* transposon libraries with both positive and negative modulation of gene expression

To characterize the functional diversity of *S. aureus*, we generated transposon libraries with similar coverage in five strains (HG003, USA300-TCH1516, MSSA476, MW2, and MRSA252). For each strain, we used our phage-based transposition method to make six sub-libraries using different transposon constructs (Fig 1A) [17]. One transposon construct contained an erythromycin resistance gene driven by its own promoter (*P_erm_*) and followed by a transcriptional terminator; another lacked the transcriptional terminator to allow readthrough from the *P_erm_* promoter; three constructs contained the transcriptional terminator, but included an additional, outward-facing promoter that varied in strength; and the sixth construct lacked the transcriptional terminator and had an added outward-facing promoter, allowing for bidirectional transcription. The sub-libraries were made separately and then pooled, with unique barcodes in each transposon construct to enable disambiguation. These high-density composite libraries often contained transposon insertions from multiple constructs at a given TA insertion site. The constructs containing outward-facing promoters minimized polar effects by allowing expression of downstream genes when an operon was disrupted. Under some conditions, they also provided supplementary fitness information. For example, under antibiotic exposure, an upregulated gene conferring a growth advantage presented in the data as a strand-biased enrichment of promoter-containing transposon insertions upstream of the gene. These “upregulation signatures” can identify molecular targets or mechanisms of resistance (Fig 1A, yellow gene; Methods).

Tn-Seq analysis showed that pooling the six transposon sub-libraries afforded libraries with high coverage in every genome (Fig 1B). Although there was a three-fold difference in unique insertions, counting insertions from each sub-library as distinct, between the library with the fewest insertions (MW2) and the library with the most insertions (USA300-TCH1516), approximately 90% of the genes in each genome had high coverage of TA dinucleotide insertion sites. This gave us confidence that we could obtain a reliable estimate of gene essentiality differences across the strains.

### Essential gene analysis reveals unexpected differences between strains

We performed Tn-Seq analysis on the five transposon libraries to identify essential genes in each strain. For the purposes of this study, we defined an essential gene as one whose loss precluded survival in competition with a heterogeneous bacterial population. We categorized genes as essential, non-essential, or indeterminate based on TA insertion site coverage and sequencing reads (S1 Table). Most essential genes were essential in all five strains. These 200 genes represent the core essential genome of *S. aureus* (S1 Fig, S2 Table), with an additional 11 that were required for at least the three MRSA strains (Fig 2A). Gene ontology overrepresentation analysis showed that the ubiquitously essential genes were enriched in DNA, RNA, and protein metabolic processes (S3 Table). Phospholipid and monosaccharide synthesis were also enriched pathways, reflecting the importance of fatty acids and other cell envelope precursors. Hypothetical genes were significantly underrepresented among the ubiquitously essential genes but dominated the group of essential genes that were only present in a subset of the genomes. Only two such genes were annotated: the antitioxin gene *pezA* found only in MW2 and a phage repressor found in HG003, MW2, and MSSA476.

**Fig 2.**
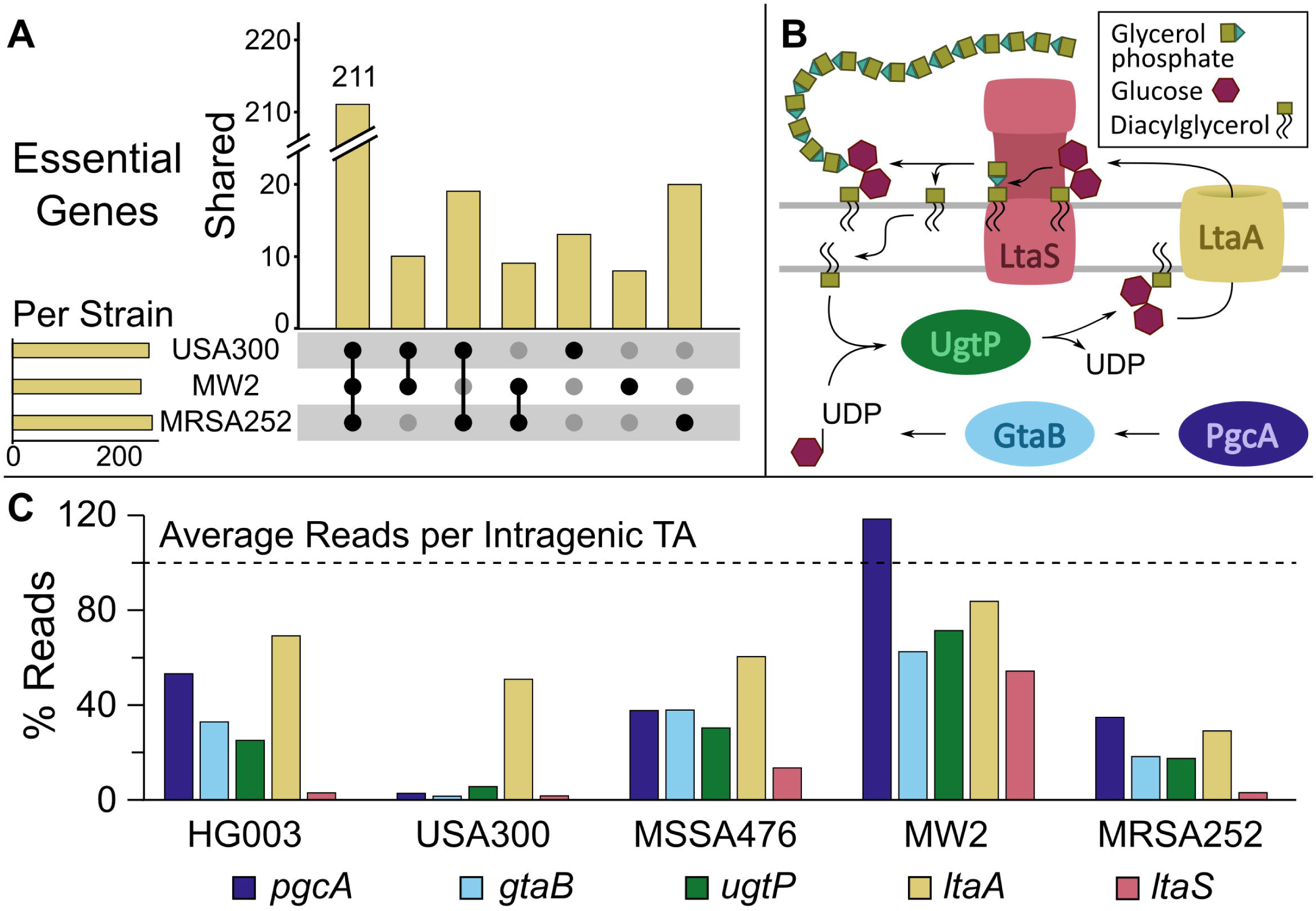
The fitness of lipoteichoic acid pathway mutants varies by strain. (A) The number of essential genes per strain (horizontal bars) and the number of genes found to be essential for all three, two, or one of the MRSA strains (vertical bars, with black dots denoting strains for which the genes are essential). Only MRSA strains were included for simplicity, but a similar plot with the MSSA strains is included in the supporting information. Note that gene essentiality is functionally defined as a lack of transposon insertions in a gene, but it may be possible to make knockouts of some of these genes under favorable conditions. (B) A schematic of the lipoteichoic acid pathway in *S. aureus*. (C) The number of reads in each LTA pathway gene for each transposon library, expressed as a percentage of the average number of reads per TA site in the coding regions of the libraries. There are strain-dependent differences in transposon insertions in LTA pathway genes, with USA300-TCH1516 appearing reliant on most genes in the pathway while MW2 is insensitive even to inactivation of *ltaS*.

Among annotated genes present in all five strains, there were notable differences in essentiality. Strains of the same sequence type displayed as much variability in gene essentiality as strains from different sequence types (S1 Fig), suggesting it may not be possible to draw general conclusions about a clonal complex from a functional genomics analysis of one member. Variably essential genes were found in multiple pathways. For example, the uniquely essential genes in MRSA252 included *sbcD*, which is homologous to an *E. coli* gene encoding a DNA hairpin cleavage enzyme [27, 28]; *sagB*, which encodes a β-N-acetylglucosaminidase [29]; and *gdpP*, which hydrolyzes the bacterial second messenger cyclic-di-AMP [30]. It is important to emphasize that essentiality in a Tn-Seq analysis does not imply that individual knockouts are not viable; however, it does suggest that there are substantial differences in the fitness of mutants across strains and that it is worth considering the broader genomic context in evaluating the cellular roles of genes.

We expected pathways involved in constructing the cell envelope to be equally essential for all strains, given the importance of the cell envelope in protecting the cell from outside stressors. We were therefore surprised to find essentiality differences within the LTA biosynthesis pathway. LTAs are long glycerol or ribitol phosphate polymers anchored in the membrane of Gram-positive bacteria. In *S. aureus*, LTAs are important in cell division, autolysin regulation, and virulence, among other processes [31]. The LTA pathway consists of five genes: *pgcA, gtaB, ugtP, ltaA*, and *ltaS* (Fig 2B). The first three genes encode proteins involved in the biosynthesis of diglucosyl-diacylglycerol (Glc_2_DAG), the membrane anchor on which LTAs are assembled. Glc_2_DAG is made inside the cell and flipped to the cell surface by LtaA. The LTA synthase, LtaS, then assembles the polymer by sequential transfer of phosphoglycerol units from phosphatidylglycerol to Glc_2_DAG. LTAs can then be further modified on the extracellular surface by D-alanine substitutions and N-acetyl glucosamine residues [32, 33]. If Glc_2_DAG is not available due to deletion of an upstream gene, LtaS can synthesize LTAs on the alternative membrane anchor phosphatidylglycerol. The four genes upstream of *ltaS* were previously found to be nonessential for *S. aureus* viability [34–36], but *ltaS* mutants were reported to be nonviable except at low temperature and under osmotically-stabilizing conditions unless suppressors were acquired [37–40]. Our Tn-Seq analysis showed, however, that mutant fitness within the LTA pathway varied considerably by strain (Fig 2C). The *ltaA* gene was expendable in all strains, but the three genes responsible for synthesis of the Glc_2_DAG anchor, *pgcA, gtaB*, and *ugtP* (*ypfP*), were uniquely essential by Tn-Seq analysis for USA300-TCH1516 (Fig 3A, S2 Fig). Conversely, *ltaS* was unexpectedly dispensable in MW2 and MSSA476 (Fig 3C, S2 Fig).

**Fig 3.**
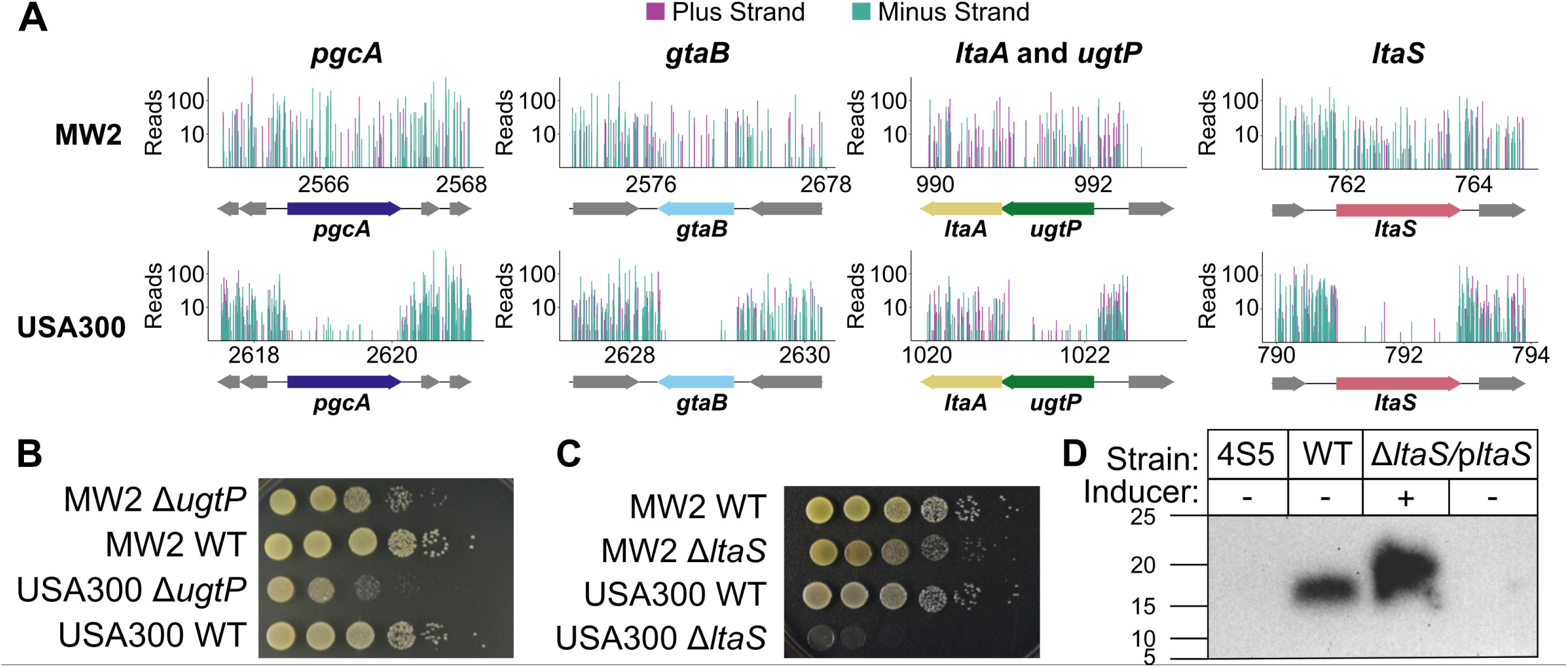
The lipoteichoic acid pathway is crucial for fitness of USA300-TCH1516 but is dispensable in MW2. (A) Tn-Seq data for the LTA pathway genes, with sequencing reads from insertions in the plus strand in purple and minus strand in teal. The reads were calculated by summing two replicate Tn-Seq experiments, only including data from transposon constructs that contained a transcriptional terminator. The data were normalized to each other by non-zero means normalization prior to plotting. The x-axis is expressed in kilobases. The y-axis is expressed on a log_10_ scale and is truncated to 500 reads. (B) Growth on agar plates for wildtype (WT) and Δ*ugtP* strains confirms that USA300-TCH1516 is more sensitive to *ugtP* deletion than MW2 is. (C) Growth on agar plates of wildtype and Δ*ltaS* strains confirms that *ltaS* is dispensable in MW2. (D) Western blot of LTAs produced by 4S5 (known to not produce LTAs), wildtype MW2, and MW2-P_Atet_-*ltaS* confirms that MW2-P_Atet_-*ltaS* does not produce LTAs unless expression of the exogenous copy of *ltaS* is induced.

We tested individual deletion mutants in the LTA pathway to confirm the Tn-Seq results. We were able to delete *ugtP* from USA300-TCH1516, consistent with results in other USA300 strain backgrounds, but a spot dilution assay showed that the mutant was less fit than the corresponding mutant in MW2 (Fig 3B). We also found that *ltaS* is dispensable in MW2, but essential in other strains (Fig 4C). Although the MW2Δ*ltaS* strain was more temperature-sensitive than MW2 wildtype, it grew on plates even at 37°C (S3 Fig). Because lethality due to *ltaS* deletion can be suppressed by deletion of *gdpP*, *clpX*, and *sgtB* [37–39], we sequenced the transposon library parent strain but found no nonsynonymous SNPs in *sgtB* or *gdpP* and only one nonsynonymous SNP in *clpX*. The amino acid difference in ClpX is unlikely to explain the observed phenotype, as it is located in an unstructured region of the protein and the amino acid is not strictly conserved in the *S. aureus* phylogeny. Moreover, we did not find any evidence for a duplication of *ltaS* in MW2 and confirmed that the MW2Δ*ltaS* mutant does not produce LTAs (Fig 3D). The explanation for why LtaS is nonessential in MW2 must lie elsewhere in the genome and will need to be investigated further.

**Fig 4.**
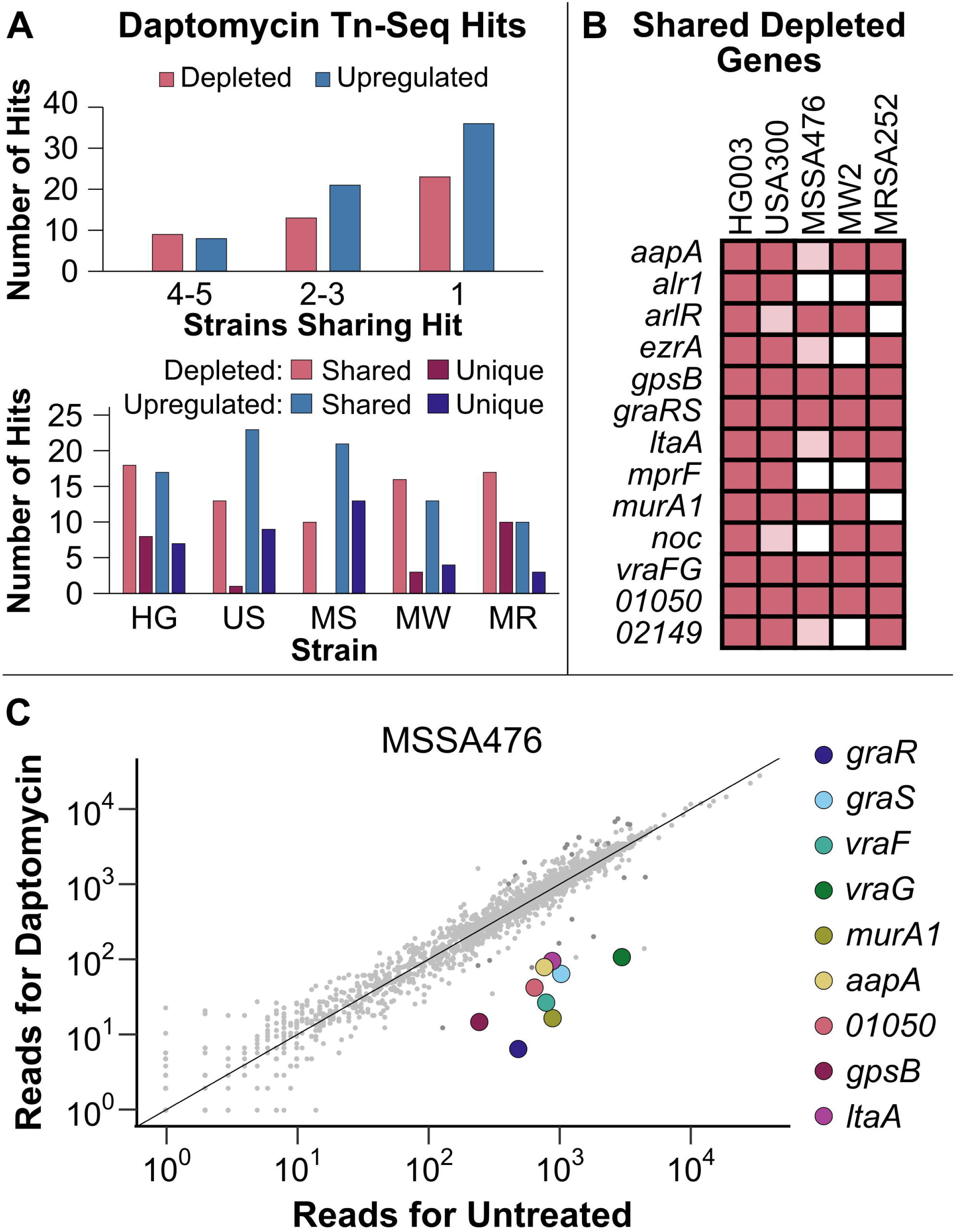
Transposon sequencing supports previously reported vulnerabilities in *S. aureus* and reveals new ones. (A) A comparison of the Tn-Seq hits under daptomycin exposure in five strains of *S. aureus*. All hits needed to have a q-value less than 0.05. Depleted genes had 10-fold fewer reads in the daptomycin-exposed sample compared to the control sample after normalization. Upregulated genes had a strand-biased enrichment of reads in the region upstream of the gene. The upper graph shows how many strains each hit was found in while the lower graph shows how many hits found in each strain were shared with at least one other strain and how many were unique to that strain. (B) Genes that were depleted of reads under daptomycin exposure in at least three of the transposon libraries, using the cutoffs described above. Hypothetical genes are listed according to their NCTC 8325 locus tag numbers. Genes meeting all cutoffs in a strain are indicated by dark pink. Those that have a significant q-value but are only depleted 5- to 10-fold are indicated in light pink. (C) Normalized Tn-Seq reads from daptomycin-exposed MSSA476 in RPMI+LB plotted against the reads from the control condition. Genes from above that were shared by at least 4 strains are highlighted.

### *S. aureus* strains share key daptomycin resistance factors

While antibiotics targeting essential genes are invaluable, it may also be advantageous to have compounds that resensitize bacteria to existing antibiotics, though they may not be antimicrobial *in vitro* on their own. To find targets for such compounds, we chose daptomycin as a case study. Daptomycin is a calcium-dependent lipopeptide antibiotic frequently used to treat *S. aureus* infections. Although its mechanism of action is not completely clear, it has been shown to insert into Gram-positive bacterial cell membranes and change membrane curvature, mislocalize cell division and cell wall synthesis proteins, depolarize the membrane, and ultimately kill the cell [41–46]. The majority of *S. aureus* infections remain susceptible to daptomycin, but there have been steady reports of daptomycin nonsusceptibility, making it imperative to identify intrinsic resistance factors that can be targeted to restore susceptibility [47].

We grew the five *S. aureus* transposon libraries in the presence and absence of daptomycin in both cation-adjusted Mueller Hinton broth (MHBII) and Roswell Park Memorial Institute supplemented with 10% lysogeny broth (RPMI+LB), as we wanted to ensure the results were not dependent on a particular medium. We found substantial overlap in the significantly depleted and upregulated genes in the two media, so the results were considered as a union of the two conditions. For every strain, there were more hits shared with at least one other strain than there were hits unique to that strain (Fig 4A, S4 and S5 Table). Genes that were substantially depleted (>10-fold) in three or more strains are shown in Fig 4B. Several vulnerabilities were above or very close to the 10-fold cutoff in all five strains.

Many of the shared vulnerabilities to daptomycin are in genes related to the cell envelope. These include *graRS/vraFG*, encoding the multi-component signaling system that regulates cell envelope processes [48–50]; *arlR*, encoding a regulator of virulence genes [51, 52]*; murA1*, encoding an enzyme that catalyzes the first step in peptidoglycan biosynthesis [53, 54]; *alr1*, one of two genes encoding an alanine racemase that supplies D-alanine for cell wall synthesis [55]; *mprF*, encoding a phosphatidylglycerol lysyltransferase that modifies membrane charge [56]; and *ltaA*, encoding the Glc_2_DAG flippase gene in the LTA pathway [34]. The earlier LTA pathway genes, *gtaB, pgcA*, and *ugtP*, were also significantly depleted in MW2, with the latter two also depleted in HG003 (Fig 5A). In addition, we identified three depleted cell division regulators, *gpsB*, *ezrA*, and *noc* (Figs 4B and 4C) [57–59]. A number of these depleted genes have been previously associated with daptomycin susceptibility, including *mprF* [60, 61], *graRS*/*vraFG* [49, 62], and *ezrA* [63], but some that were substantially depleted in all five strains have not been connected to daptomycin, including *gpsB* and *ltaA*. We confirmed that deletion of *ltaA* sensitizes HG003, MW2, and USA300-TCH1516 to daptomycin and that the sensitivity can be reversed by *ltaA* complementation (Fig 5B, S4 Fig). Therefore, Glc_2_DAG-LTA in the cell membrane contributes to *S. aureus* survival in the presence of daptomycin, and it may be possible to exploit this observation to overcome daptomycin nonsusceptibility or limit its development.

**Fig 5.**
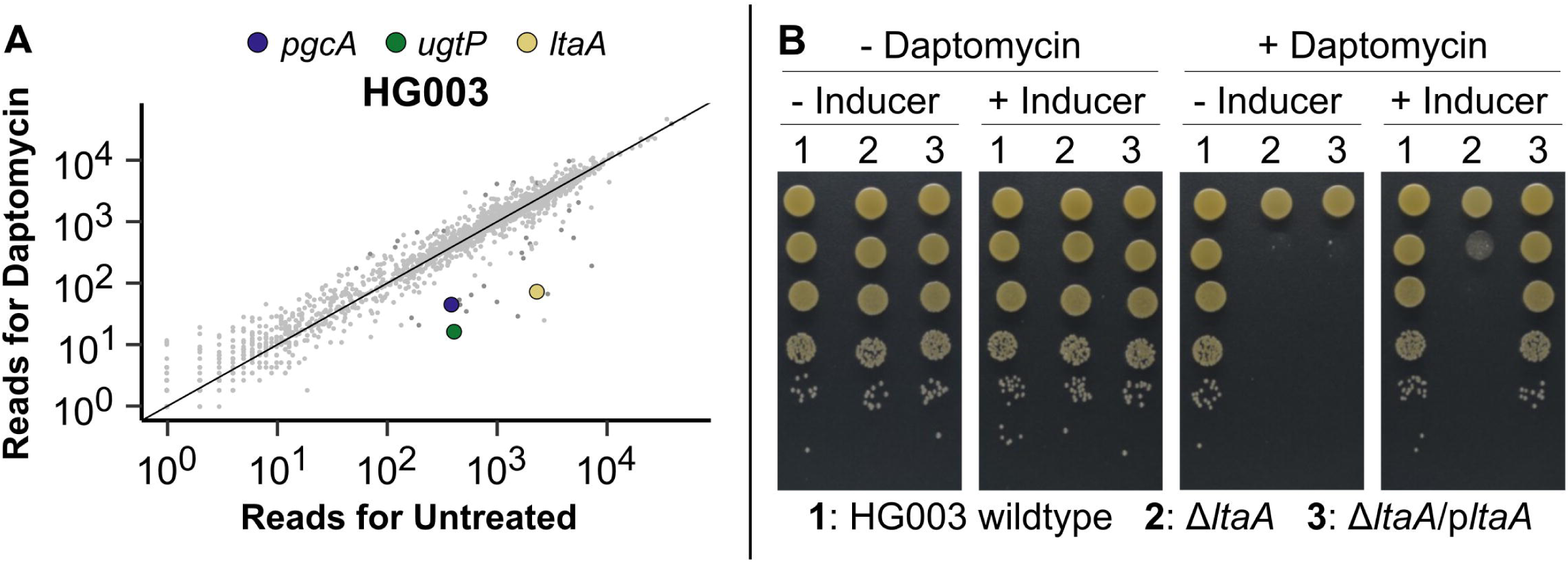
LTA loss sensitizes *S. aureus* to daptomycin. (A) Normalized Tn-Seq reads from daptomycin-exposed HG003 in RPMI+LB plotted against the reads from untreated HG003 in the same growth medium. Genes in the LTA pathway that were significantly depleted are highlighted. (B) Growth on agar plates of HG003 wildtype, Δ*ltaA*, and inducible *ltaA* complementation strains in the presence or absence of daptomycin and the inducer confirms that *ltaA* is required for *S. aureus* growth under daptomycin exposure.

Just as there were more shared than unique depleted genes for each strain, there were more shared than unique upregulated genes (Fig 4A). The outward-facing promoters in several of our transposon constructs can upregulate downstream genes, allowing us to identify genes that confer a fitness advantage when upregulated. They are detected by strand-biased read enrichments in promoter-containing constructs, and we have previously confirmed that these strand-biased read enrichments, or upregulation signatures, do reflect increased expression of proximal genes [19]. Often the genes identified this way are the targets of the antibiotic itself or reveal gain-of-function resistance mechanisms. Standard transposon libraries cannot identify gain-of-function resistance mechanisms, although loss-of-function resistance mechanisms can be identified through intragenic read enrichments. Here we found that the genes enriched in reads upon daptomycin exposure belonged to a wide assortment of cellular processes and we could not discern a pattern amongst them (S6 Table).

Many of the genes for which we identified upregulation signatures in multiple strains were previously implicated in daptomycin resistance in *S. aureus* (Table 1). Daptomycin nonsusceptibility normally develops through the acquisition of multiple independent mutations that result in changes to the cell envelope [47], and many of the mutations are thought to be gain-of-function mutations that result in increased activity. For example, mutations in *mprF* have been selected *in vitro* and in the clinic and are thought to increase lysyl phosphatidylglycerol in the outer leaflet of the membrane [64]. Other gain-of-function resistance mechanisms identified in our Tn-Seq libraries include *cls2*, one of two cardiolipin synthases in *S. aureus* [65]; the staphyloxanthin biosynthesis gene *crtM* [66]; and two signaling systems, *graRS* and *walKR* (*yycFG*) [62, 67]. Some of these resistance mechanisms have been reported for daptomycin resistance in other bacteria as well [68]. These hits affirm the utility of upregulation signatures for identifying clinically-relevant mechanisms of resistance. The gene *murA2*, whose function was identified as important for withstanding daptomycin exposure in the depletion analysis, also had upregulation signatures in several strains. The importance of MurA activity identified in this study – both through the depletion analysis and the upregulation analysis – is consistent with recent studies showing that a combination of daptomycin and fosfomycin is significantly more effective at curing MRSA bacteremia than daptomycin alone, leading to an ongoing phase III clinical trial [69–71]. One caveat of the upregulating transposons is that they may have distal effects, upregulating both the most proximal gene and other downstream genes, especially within operons. Individual mutants need to be tested to confirm that the proximal gene is most responsible for the phenotype. For example, we identified an upregulation signature for *ugtP* in USA300-TCH1516 and in MSSA476. The *ugtP* gene is upstream of *ltaA* in an operon, so it is unclear whether upregulating *ugtP* on its own would recapitulate the transposon-induced fitness advantage or whether *ltaA* upregulation is also necessary. Regardless of whether upregulation of one or both genes in the operon drives the fitness advantage, the result is consistent with findings from the depletion analysis that Glc_2_DAG-LTA is important for withstanding daptomycin stress.

**Table 1.**
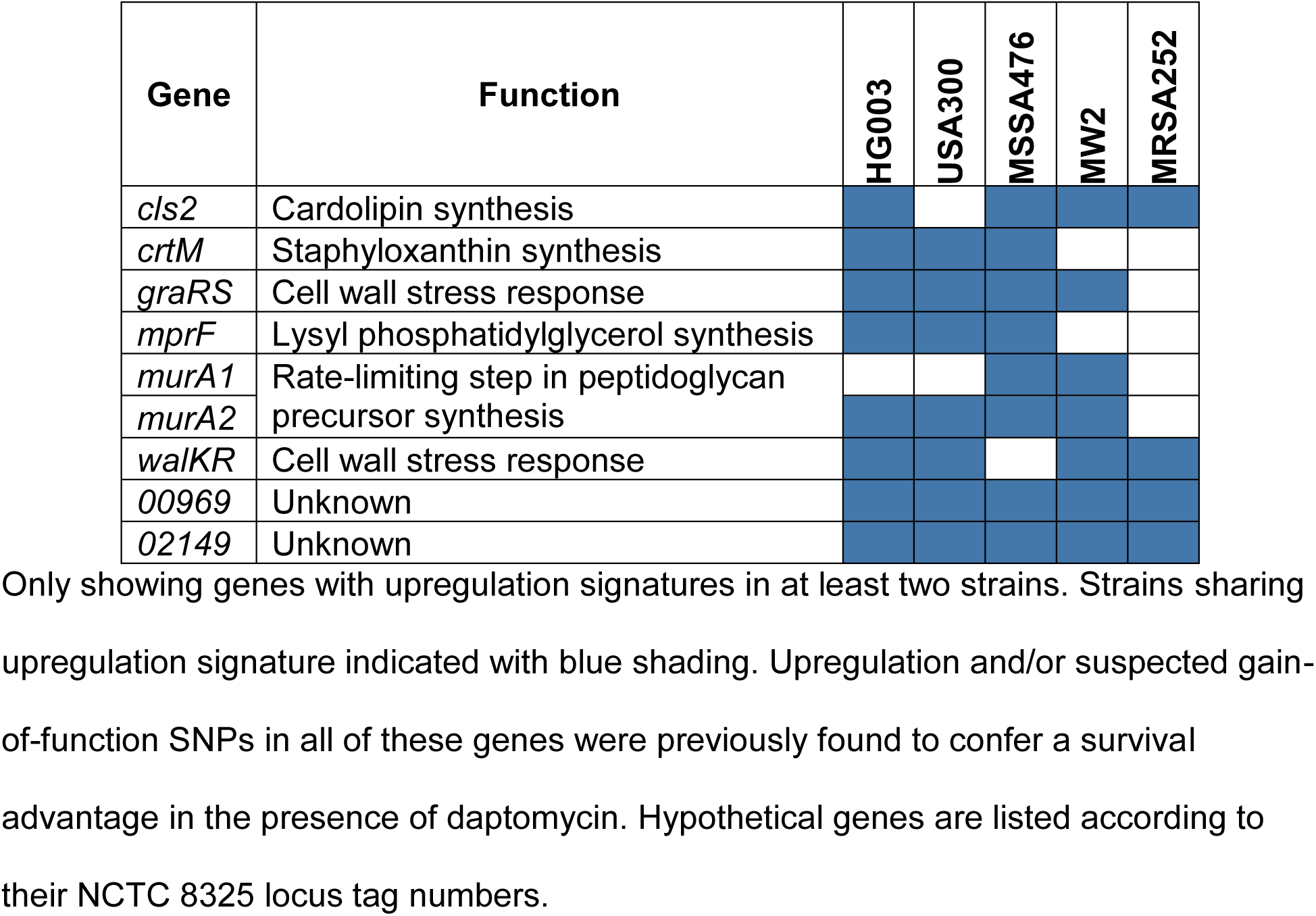
Upregulation signatures identify genes previously linked to reduced susceptibility to daptomycin.

Upregulation signatures were found upstream of the genes *SAOUHSC_02149* and *SAOUHSC_00969* in all five strains. These genes each encode a small protein of unknown function with a single transmembrane helix. We previously identified both as daptomycin resistance factors in a Tn-Seq study of a single *S. aureus* strain and confirmed that upregulation confers resistance while inactivation increases susceptibility to daptomycin [19]. The observation that these genes are important for daptomycin resistance in all five *S. aureus* strains suggests some effort should be devoted to evaluating the physiological roles of their encoded products.

## Discussion

Declining sequencing costs and efficient transposon mutagenesis methodologies enable investigation of the genetic basis of phenotypes on a large scale. Tn-Seq experiments are typically conducted in a single strain that is considered representative, and it is assumed that results from that strain can be extrapolated to other members of the species. Recent studies have challenged this notion, demonstrating clear strain-dependent gene requirements in *Mycobacterium tuberculosis, Streptococcus pneumoniae*, and *Pseudomonas aeruginosa* [18, 26, 72].

In the present study, we reveal how complicated the concept of gene essentiality can be. We found that the LTA pathway, which has always been presumed to be essential for all *S. aureus* strains, was variably essential under normal growth conditions, ranging from required in USA300-TCH1516 to expendable in MW2. We suspect that intracellular Glc_2_DAG has an unidentified function that USA300-TCH1516 depends upon for survival because the Glc_2_DAG biosynthesis genes (*pgcA, gtaB*, and *ugtP)* contained almost no transposon reads, whereas *ltaA*, which encodes the flippase responsible for translocating Glc_2_DAG to the cell surface, contained many insertions.

The central role of Glc_2_DAG in the viability of USA300-TCH1516 may provide a tool to interrogate what other roles the molecule has in the cell. Likewise, having a strain like MW2 that can ostensibly survive without LTAs provides new opportunities for characterizing the functions of LTAs, as future investigations can interrogate what wildtype MW2 can do that an *ltaS* knockout cannot. Whether strains that can survive without LTA *in vitro* – due to the presence of known suppressors or to other uncharacterized genetic background differences – can survive in the more challenging environment of an animal infection needs to be assessed. LTAs are associated with virulence and it has been shown in the *S. aureus* Newman strain that *ltaA* is required for fitness in a mouse abscess model [34]. The results here also show that the LTA pathway becomes universally required in the presence of daptomycin; therefore, LTA pathway inhibitors may resensitize nonsusceptible isolates to daptomycin even if they cannot kill the cells on their own.

We have also demonstrated an improved method for identifying gain-of-function resistance mechanisms via Tn-Seq. The upregulation signatures that we found under daptomycin exposure using our transposons with outward-facing promoters largely echoed the depletion hits. In other words, if removing a gene sensitizes a bacterium to an antibiotic, upregulating that gene is often protective. For daptomycin, these intrinsic resistance factors were predominantly cell envelope, cell division, and stress response system genes. Although a standard Tn-Seq library can identify intrinsic resistance factors through sensitization of knockouts, sensitization readouts are limited to only looking at nonessential genes where a decline in insertions can be detected. The upregulation signatures, on the other hand, are indifferent to the essentiality of the gene. For example, *ugtP* was upregulated under daptomycin stress in USA300-TCH1516 even though it was found to be essential in that strain, and *walR*, an essential multicomponent signaling system gene, was upregulated in multiple strains. As validation for the approach, most of the genes that had upregulation signatures have been previously reported as suspected gain-of-function resistance mechanisms for daptomycin.

In summary, our comparison of Tn-Seq data across multiple strains provided a more nuanced illustration of *S. aureus* genetic dependencies than we could have attained using a single strain. The comparative data provide the most comprehensive list of core essential genes in *S. aureus* to date and have identified myriad strain-dependent essential genes. The strain-to-strain gene essentiality differences under normal growth conditions provide a basis to more deeply interrogate *S. aureus* biology. Meanwhile, those factors that were universally essential may be useful targets for future antibiotic development. Under daptomycin stress the commonalities between the strains likewise proved valuable, as they give us a means to prioritize targets for the development of synergistic compounds. Moreover, Tn-Seq results can be used to complement findings from other systems biology approaches [9, 73]. As sequencing continues to become more affordable, we envision performing Tn-Seq in multiple strains will become routine and will lead to a more comprehensive understanding of the scope of bacterial phenotypic diversity.

## Materials and methods

### Materials, bacterial strains, plasmids, and oligonucleotides

All reagents were purchased from Sigma-Aldrich unless otherwise indicated. Bacterial culture media were purchased from BD Sciences. Restriction enzymes and enzymes for Tn-seq preparation were purchased from New England Biolabs. Oligonucleotides and primers were purchased from Integrated DNA Technologies. DNA concentrations were measured using a NanoDrop One Microvolume UV-Vis Spectrophotometer (Thermo Scientific). DNA sequencing was performed by Eton Bioscience unless otherwise noted. KOD Hotstart DNA polymerase (Novagen) was used for PCR amplification. *E. coli* strains were grown with shaking at 37°C in lysogeny broth (LB) or on LB plates. *S. aureus* strains were grown at 30°C in tryptic soy broth (TSB) shaking or on TSB agar plates unless otherwise noted. For *S. aureus* strains, compounds for selection or gene induction were used at the following concentrations: 5 µg/mL chloramphenicol and erythromycin; 50 μg/mL kanamycin and neomycin; or 2.5 μg/mL tetracycline. For *E. coli* strains, 100 µg/mL carbenicillin was used for selection. The bacterial strains, plasmids, and oligonucleotide primers used in this study are summarized in S7 Table.

### Transposon library construction

Construction of the transposon library in the laboratory *S. aureus* strain HG003 has been described [17]. A similar strategy was used to make libraries in other *S. aureus* genetic backgrounds after deleting non-compatible DNA restriction systems and endogenous antibiotic resistance genes that would interfere with transposon library construction. In the community acquired *S. aureus* MW2 and MSSA476 strains, the clonal complex CC1 *hsdR* Type I restriction system in each background was first deleted using the temperature sensitive shuttle vector pKFC with ~1 kb DNA homology flanking arms. A second restriction system unique to MSSA476 was deleted in a subsequent round using pKFC to generate the transposon library host. Libraries were constructed in the *S. aureus* community acquired strain USA300-TCH1516 by first curing the endogenous plasmid pUSA300HOUMR through destabilizing replication via integration of pTM283. Cointegrated pUSA300HOUMR-pTM283 was passaged at 30°C for two rounds of outgrowth in 10 mL of TSB media before streaking to single colonies. Colonies exhibiting kanamycin and erythromycin sensitivity (encoded on pUSA300HOUMR) were further checked by PCR to confirm loss of plasmid. The resulting strain, TM283, was used as host for transposon library construction. A transposon library of the hospital-acquired, methicillin-resistant *S. aureus* MSRA252 strain was made by first converting the *hsdR* (SAR0196) Type I restriction gene deletion shuttle vector pGKM305 into a high frequency transduction vector by adding a φ11 DNA homology region (primers GKM422-423) into the SfoI site to make pGKM306. The plasmid was electroporated into RN4220 and transduced into wildtype MRSA252 using φ11-FRT with temperature permissive selection at 30°C [17]. The SAR0196 gene was then deleted as described above. Both copies of the endogenous duplicated erythromycin resistance gene (SAR0050 and SAR1735) were likewise deleted in two rounds using pKFC_SAR0050, except plasmids were directly electroporated into the restriction negative parent strain GKM361. Due to endogenous aminoglycoside resistance, the transposase expressing plasmid pORF5 Tnp+ was converted into chloramphenicol resistant plasmids to enable selection in strain TXM369 and library construction.

### Transposon sequencing

To identify the essential genes in each of the *S. aureus* strains (HG003, USA300-TCH1516, MSSA476, MW2, and MRSA252), library aliquots were thawed and diluted to an OD_600_ between 0.2 and 0.3 in 10 mL of MHBII in duplicate. The cultures were then incubated shaking at 30°C until they reached an OD_600_ of approximately 0.4, roughly 1.5-1.75 hours. The cells were pelleted, and the DNA was extracted and prepared for Tn-Seq as previously described [17]. Samples were then submitted to either the Harvard Biopolymers Facility or the Tufts University Core Facility for sequencing on a HiSeq 2500 instrument.

To identify genes affected by daptomycin exposure, transposon library aliquots for the six strains were inoculated in either MHBII or RPMI+LB. The cultures were incubated for 1.5 hours at 30°C in a shaking incubator as described above. The cultures were diluted to OD_600_ 0.005 in either 2 mL MHBII or RPMI+LB containing a series of daptomycin concentrations (μg/mL): 0, 0.12, 0.25, 0.5, and 1. The cultures were then incubated at 37°C in a shaking incubator and their ODs were monitored using a Genesys 20 spectrophotometer (Thermo Scientific). The 2 mL cultures were harvested by centrifugation when the OD_600_ reached 1.5. The cell pellets were stored at −80°C until processing for Tn-Seq. The genomic DNA was extracted and prepared as described previously [17]. Samples were submitted to the Tufts University Core Facility for sequencing on a HiSeq 2500 instrument.

### Transposon sequencing data analysis

Transposon sequencing data was split by transposon and sample, trimmed, filtered, and mapped using the Galaxy software suite as previously described [17, 19, 74, 75]. A workflow for the processing is provided on GitHub. The resulting SAM files were converted into tab-delimited hop count files using Tufts Galaxy Tn-Seq software (http://galaxy.med.tufts.edu/) or custom python scripts, likewise provided on GitHub, and then converted further into IGV-formatted files, as previously described [17, 19].

Chromosome nucleotide FASTA files for NCTC 8325 (NC_007795.1 - HG003 parent strain, as the HG003 genome is not closed), USA300-TCH1516 (NC_010079.1), MSSA476 (NC_002953.3), MW2 (NC_003929.1), and MRSA252 (NC_002952.2) were downloaded from the NCBI genomes database. The genomes were reannotated via Prokka and the pangenome was aligned with Roary, splitting by homolog and using a 90% ID cutoff [76, 77]. Roary group names were then adjusted based on common *S. aureus* pangenome gene names found on AureoWiki [78].

Genes in each strain were labeled as essential, non-essential, or uncertain using the TRANSIT software Gumbel method [79]. Only the transposon constructs with transcriptional terminators were included (four of six transposon constructs). For each TRANSIT Gumbel run, data files for each transposon construct in both replicates (for a total of eight files) were submitted and the mean replicates parameter was chosen. Permutation tests with 20,000 permutations were then conducted between each pair of strains for each gene to determine whether differences in essentiality were significant, using the sum of the same data sets used in the Gumbel analysis. Data were normalized by average reads per TA site with reads (non-zero means normalization) and the p-values from each file pair were corrected for multiple hypothesis testing using the Benjamini-Hochberg method. A gene was considered significantly different between two strains if the q-value was less than 0.05. The list of genes essential for all five strains was then analyzed using the online gene ontology tool PANTHER version 13.1, comparing the NCTC 8325 locus tags for the essential genes to the *S. aureus* reference gene list and using default settings for the overrepresentation test [80].

Genes depleted or enriched under daptomycin stress were identified by normalizing the treated sample data by non-zero means and performing a Mann-Whitney U test for each gene. The p-values were then corrected for multiple hypothesis testing by the Benjamini-Hochberg method. A depleted gene was considered to be a hit if there were at least 100 reads in the control file, the q-value was less than 0.05, and the treated:control read ratio was less than 0.1. An enriched gene was considered to be a hit if there were at least 100 normalized reads in the treated file, the q-value was less than 0.05, and the read ratio was greater than ten. We then selected one representative daptomycin concentration for each strain in each medium, chosen to have a similar selective pressure (Table 2). Specifically, the highest concentration file with twenty-five or fewer 10-fold depleted genes with a significant q-value was chosen.

**Table 2.** Daptomycin samples included in the multi-strain comparison of depleted and upregulated genes.

| Strain | Medium | Concentration (µg/mL) | Depleted Genes |
| --- | --- | --- | --- |
| USA300-TCH1516 | RPMI+LB | 0.12 | 8 |
| USA300-TCH1516 | MHBII | 0.5 | 12 |
| MW2 | RPMI+LB | 0.25 | 18 |
| MW2 | MHBII | 0.5 | 8 |
| MSSA476 | RPMI+LB | 0.25 | 7 |
| MSSA476 | MHBII | 0.5 | 8 |
| HG003 | RPMI+LB | 0.25 | 9 |
| HG003 | MHBII | 1 | 25 |
| MRSA252 | RPMI+LB | 0.5 | 19 |
| MRSA252 | MHBII | 1 | 18 |

Genes upregulated under daptomycin stress were identified by a bootstrapping approach. We defined the upstream region (UR) of a gene to be the 500 bp ahead of a gene, based on the 99^th^ percentile of 5’ untranslated regions of transcripts in *E. coli* [81]. Only TA sites located in a UR that had at least one read in either the control or the treated data were included in the analysis. Read differences for each site were then calculated by subtracting the control data reads from the treated data reads. For each strand and for each gene, a distribution for the expected difference in UR reads between the treated sample and the control was created by counting the number of qualifying TA sites in the UR (N) and generating 200,000 summed random samples of N read differences from across the genome. A one-sided p-value for each strand was calculated as the proportion of the null distribution values that was greater than the actual value for that gene, and a q-value was obtained using the Benjamini-Hochberg method. We defined an upregulation signature as any UR in which the q-value for the DNA strand matching the gene direction was less than 0.05, there was more than one TA site in the UR, and the opposing strand had a read difference less than the 90^th^ percentile for all URs. Again, only those samples shown in Table 2 were included.

### Whole genome sequence comparisons

To obtain the genomic sequences of the transposon library parent strains, the strains were cultured in MHBII at 37°C shaking overnight. The DNA was harvested using a Promega Wizard Genomic DNA purification kit and cleaned using a Zymo DNA Clean Up Kit. DNA was tagmented using the Illumina Nextera DNA Library Prep kit, but using 1/20^th^ of the volume recommended by the manufacturer and starting DNA concentrations of 0.5, 0.75, 1, and 2 ng/μL. The tagged DNA fragments were then amplified via PCR. The PCR samples contained 11.2 μL of KAPA polymerase mix (Illumina), 4.4 μL each of the 5 μM column and row indexing primers, and 2.5 μL of tagmented DNA. The thermocycler settings were as follows: preincubation (3 min, 72 °C), polymerase activation (5 min, 98 °C), 13 amplification cycles (denaturation at 98 °C for 10 sec, annealing at 62°C for 30 sec, and extension at 72°C for 30 sec), and termination (5 min, 72 °C).

To determine which starting DNA concentrations yielded the best fragment size, 3.75 μL of each amplified sample was mixed with 4 μL of 6x loading dye and run on a 1.5% agarose gel at 110 V. The sample with an average length closest to 500 bp was chosen for each strain. Those samples were then cleaned as recommended by Illumina, except that we started with 15 uL of amplified tagmented DNA and 12 μL of AMPure XP beads. The concentrations of the resulting DNA samples were estimated via Qubit Fluorometric Quantification (Thermo Fisher Scientific) following the manufacturer’s instructions. The DNA was then diluted to 1 ng/μL and pooled, and a portion of it was submitted to the Harvard Biopolymer’s Facility for TapeStation and qPCR quality control analysis. Upon passing the quality control step, the DNA was prepared and loaded into the sequencing cartridge as directed by the MiSeq Reagent Kit v3 with 150 cycles (Illumina), including a 1% PhiX control spike-in prepared according to manufacturer’s instructions (PhiX Control v3, Illumina), and paired-end sequenced using a MiSeq instrument.

Genomic data was then analyzed to find SNPs, insertions, and deletions. MetaPhlAn2 was used to verify that the DNA was not contaminated. Sequences were then aligned to the appropriate NCBI reference genome using the Burrow-Wheels Aligner (BWA-MEM, v7.12) with default settings [82]. Note that NCTC 8325 was used for HG003, as the HG003 genome on NCBI is not closed. Duplicate reads were marked with Picard, and Pilon (v1.16) was used for variant calling, with a minimum depth of 1/10^th^ of the average read depth and a minimum mapping quality of 30, unless the average read depth was less than 100, in which case the minimum mapping quality was set to 15 [83, 84]. The resulting VCF files were then evaluated for mutations in relevant areas of the genome (*i.e.* the LTA pathway or known *ltaS* suppressors). We compared the predicted protein sequences for LTA pathway members and *ltaS* suppressors in the five strains using the online Clustal Omega tool from EMBL-EBI [85].

### Gene deletion and complementation

To make an anhydrotetracycline (Atet) inducible construct of *ugtP*, a fragment containing the *ugtP* and *ltaA* operon (SAOUHSC_00953-00952) and its ribosomal binding site was amplified from HG003 genomic DNA using *ugtP*-F and *ugtP*-R. The fragment was then cloned into pTP63 using KpnI and EcoRI to generate pTP63-*ugtP*, and the pTP63-*ugtP* was transformed into a wildtype RN4220 strain containing pTP44 [86]. Next, the integrated inducible *ugtP* operon was transduced into wildtype HG003, MW2, and USA300-TCH1516 using φ11 phage to generate HG003-P_Atet_-*ugtP*, MW2-P_Atet_-*ugtP*, and USA300-TCH1516-P_Atet_-*ugtP*. To generate inducible *ugtP* strains, the wildtype P_Atet_-*ugtP* strains were grown in TSB containing 0.3 μM Atet for 6 hours at 30°C, after which they were transduced with a transposon-inactivated *ugtP* marked with an erythromycin resistance gene [13]. The desired mutants were selected on TSB agar containing 5 μg/mL erythromycin and 0.3 μM Atet and confirmed by PCR using the primers *ugtP*-CA and *ugtP*-CB.

To construct an Atet-inducible *ltaA* construct, the *ltaA* gene (*SAOUHSC_00952*) and the *ugtP-ltaA* operon ribosome binding site were amplified from HG003 genomic DNA using primers *ltaA*-F and *ugtP*-R and cloned into pTP63 to generate pTP63-*ltaA*. As described above, the pTP63-*ltaA* was transduced into wildtype HG003 and MW2, and the resulting strains were then transduced with a Δ*ltaA* construct marked with a kanamycin resistance gene to generate the inducible *ltaA* strains [87].

To construct an Atet-inducible *ltaS* construct, the *ltaS* gene and its ribosome binding site were PCR amplified using the primers *ltaS*-F and *ltaS*-R and cloned into pTP63 to make pTP63-*ltaS*. To make an inducible *ltaS* strain, pTP63-*ltaS* and then a Δ*ltaS* construct marked with an erythromycin resistance gene were transduced into wildtype HG003, MW2, and USA300-TCH1516 as described above [37]. The *ltaS* mutants were confirmed by PCR using the primers *ltaS*-CA and *ltaS*-CB.

### Spot dilution assays

Overnight cultures of the relevant strains were grown in TSB at 30°C until mid-log phase and diluted to an OD_600_ of 0.1. Five 10-fold dilutions of the resulting cultures were prepared, and 5 μL of each dilution was spotted on TSB plates with or without 0.3 μM Atet and, where indicated, 2.5 μg/mL daptomycin. Plates were imaged after approximately 16 hours of incubation. For the *ltaS* experiments, plates were incubated at 30°C, 37°C, and 42°C. For the *ltaA* experiments, plates were incubated at 37°C.

### LTA western blot

LTAs from MW2, MW2-P_Atet_-*ltaS*, and 4S5, a strain derived from RN4220 known to not produce LTAs [37], were isolated and detected via western blot using a procedure similar to that previously described [88]. Overnight cultures of each strain were grown shaking at 30°C in TSB supplemented with 7.5% salt, with the MW2-P_Atet_-*ltaS* grown both in the presence and absence of 0.4 μM Atet. Cultures were then diluted 1:50 in the same medium and incubated shaking at 30°C until the OD_600_ was between 0.6 and 0.75. The equivalent of 1 mL of OD_600_ 0.8 was then harvested from each sample by centrifuging at 8000 x *g* for three minutes. The cell pellets were resuspended in 50 μL of a buffer composed of 50 mM Tris pH 7.5, 150 mM NaCl, and 200 μg/mL lysostaphin. Samples were incubated at 37°C for ten minutes. Then, 50 μL of 4x SDS-PAGE loading buffer was added and the samples were boiled for thirty minutes. After returning the samples to room temperature, 100 μL water and 0.5 μL of proteinase K (New England Biolabs) was added to each sample, and samples were then incubated at 50°C for two hours. After cooling samples to room temperature, 2 μL of AEBSF was added to each sample to quench the proteinase K. Ten microliters of each sample and 5 μL of Precision Plus Protein Dual Xtra standard (BioRad) were loaded onto a 4-20% TGX precast gel (BioRad) and the gel was run for 30 minutes at 200 V using a running buffer that was 0.5% Tris, 1.5% glycine, and 0.1% SDS. Proteins were transferred to a methanol-activated PVDF membrane using a TransBlot Turbo (BioRad) via the pre-installed mixed molecular weight setting. The membrane was rinsed with TBST (0.05% Tween-20) and incubated rocking overnight at 4°C in 5% milk in TBST to block. After washing the membrane with TBST three times for five minutes each, the LTAs were bound with a 1:750 solution of a mouse anti-LTA antibody (Hycult Biotech) in TBST for 45 min. The membrane was then washed an additional three times for five minutes each before incubating in a 1:2000 dilution of anti-mouse horseradish peroxidase conjugated antibody (Cell Signaling Technologies) in TBST. The membrane was washed a final five times for five minutes each and then exposed to ECL western blotting substrate (Pierce). The membrane was imaged on a FluoroChem R gel doc (ProteinSimple) with a 2 min 40 second exposure for luminescence to detect the LTAs and automatic exposure for red and green fluorescence channels to detect the protein standard.

## Acknowledgements

We thank David Hooper (Massachusetts General Hospital) for providing the MW2 strain, Samir Moussa (Harvard Medical School) for creating the pWV01-C9 plasmid with chloramphenicol resistance, Mohamad Sater (Harvard T.H. Chan School of Public Health) for guidance in aligning and annotating whole genome sequences, and Scott Olesen (Harvard T.H. Chan School of Public Health) for productive conversations regarding the statistical analysis of transposon sequencing data.

## Supporting information

**S1 Fig. Gene essentiality varies by strain.** The number of essential genes per strain (horizontal bars) and the number of genes found to be essential for all five or a subset of the *S. aureus* strains (vertical bars, with black dots denoting strains for which the gene is essential).

**S2 Fig. Tn-Seq data reveals variable dependence on the lipoteichoic acid pathway among *S. aureus* strains.** Each column of graphs represents a gene in the LTA pathway, with rows representing strains. Reads in the Tn-Seq data, expressed on a log_10_ scale, are plotted against the position in the genome indicated on the x-axis with a depiction of the gene context underneath each plot. The y-axis is truncated to 500 reads. Purple lines indicate plus-strand reads. Teal lines indicate minus-strand reads.

**S3 Fig. MW2Δ*ltaS* mutants are temperature sensitive.** Growth on agar plates of wildtype and *ltaS* complementation strains for HG003, MW2, and USA300-TCH1516 with and without inducer at 30°C, 37°C, and 42°C.

**S4 Fig. *ltaA* is conditionally essential in USA300-TCH1516 and MW2 in the presence of daptomycin.** Growth on agar plates of wildtype, Δ*ltaA*, and inducible *ltaA* complementation strains for MW2 and USA300-TCH1516 in the presence or absence of daptomycin and the complementation inducer confirms that *ltaA* is required for *S. aureus* growth under daptomycin exposure.

**S1 Table. Comparison of essential genes in *S. aureus* strains.** A table of all genes that are essential for at least one strain, with essentiality designations for each gene in each strain for the TRANSIT Gumbel analysis and the subsequent permutation test which was used to determine whether differences between strains seen in the Gumbel results were significant. If differences were not significant, essential (E) and nonessential (NE) designations were converted to uncertain (U).

**S2 Table. Core essential genes in *S. aureus*.** A list of the genes that were determined to be essential in all five *S. aureus* strains.

**S3 Table. Gene ontology analysis reveals central dogma pathways to be universally essential in *S. aureus*.** Red text highlights those processes mentioned in the main text. DNA, RNA, and protein metabolic processes are significantly over-represented among the genes essential for all five strains of *S. aureus* studied. Phospholipid and monosaccharide synthesis were likewise over-represented, while genes of unknown function were under-represented.

**S4 Table. Genes depleted under daptomycin exposure.** A table of the genes that were 10-fold depleted of reads in the normalized daptomycin file compared to the control file with q-values less than 0.05 and at least 100 reads in the control file.

**S5 Table. Genes upregulated under daptomycin exposure.** A table of the genes that had upregulation signatures under daptomycin exposure. See methods for details of analysis.

**S6 Table. Genes enriched under daptomycin exposure.** A table of the genes that were 10-fold enriched in reads in the normalized daptomycin file compared to the control file with q-values less than 0.05 and at least 100 normalized reads in the daptomycin file.

**S7 Table. Primers, plasmids, and strains used in this study.**

