## Supplementary figures and images for "Comparative Tn-Seq reveals common daptomycin resistance determinants in *Staphylococcus aureus* despite strain-dependent differences in essentiality of shared cell envelope genes"

### S1 Fig

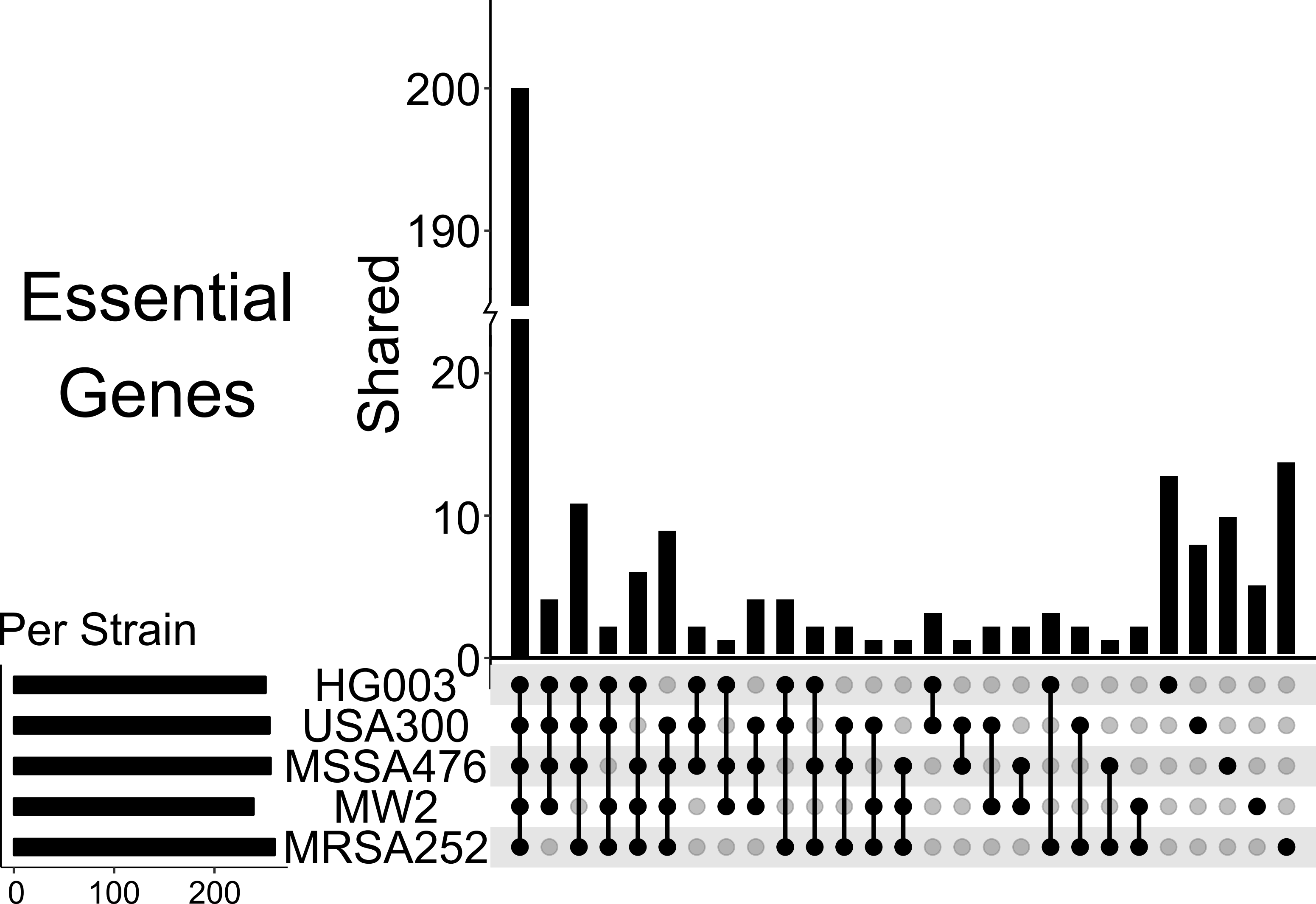

### S2 Fig

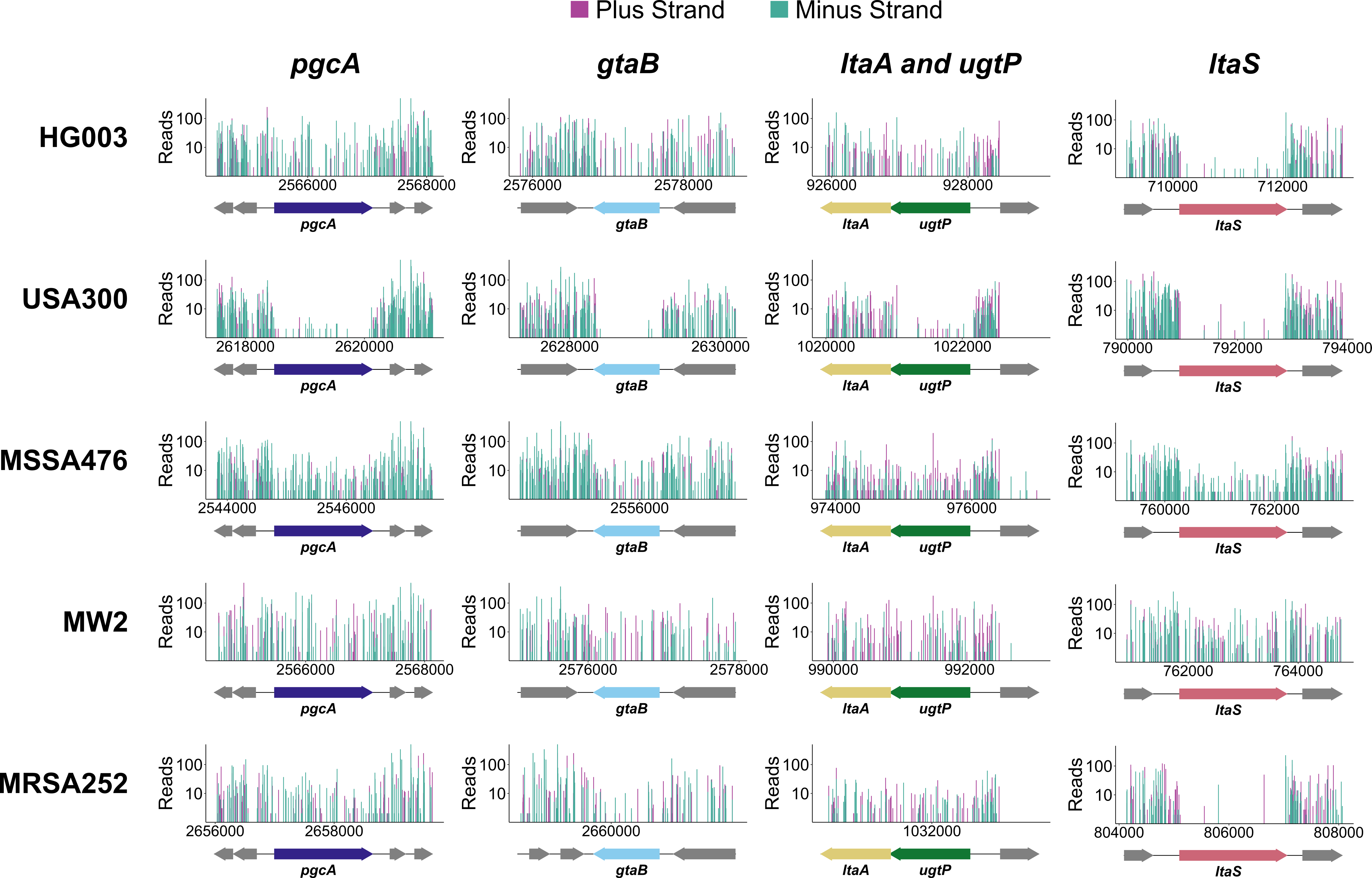

### S3 Fig

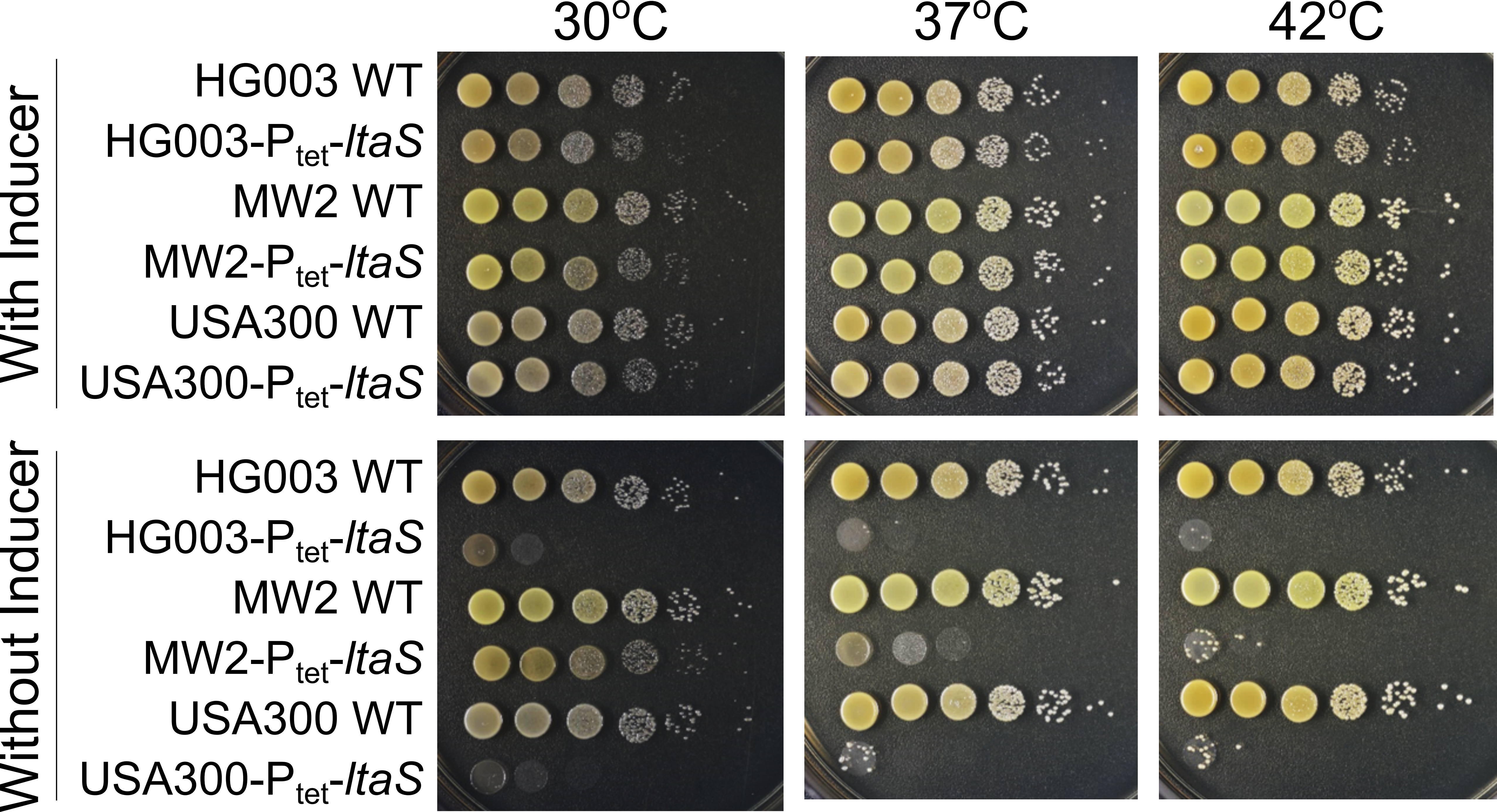

### S4 Fig

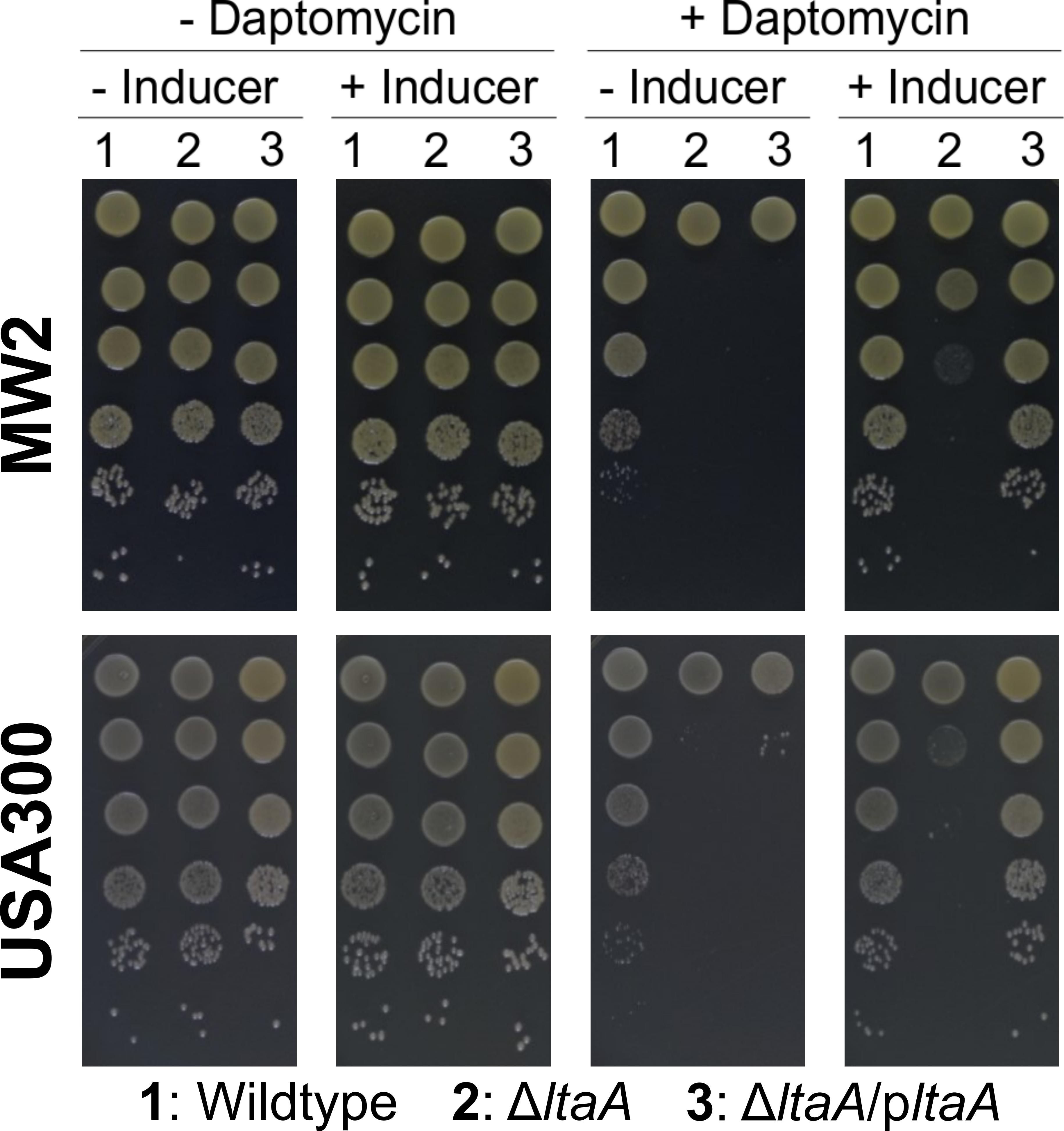
